# Cross-neutralization antibodies against SARS-CoV-2 and RBD mutations from convalescent patient antibody libraries

**DOI:** 10.1101/2020.06.06.137513

**Authors:** Yan Lou, Wenxiang Zhao, Haitao Wei, Min Chu, Ruihua Chao, Hangping Yao, Junwei Su, Yanan Li, Xiulan Li, Yu Cao, Yanyan Feng, Ping Wang, Yongyang Xia, Yushuan Shang, Fengping Li, Pingju Ge, Xinglin Zhang, Wenjing Gao, Bing Du, Tingbo Liang, Yunqing Qiu, Mingyao Liu

**Affiliations:** State Key Laboratory for diagnosis and treatment of infectious diseases, Key Laboratory for Drug Evaluation and Clinical Research of Zhejiang Province, The First Affiliated Hospital, Zhejiang University School of Medicine, Hangzhou 310003, China; Shanghai Key Laboratory of Regulatory Biology, Institute of Biomedical Sciences and School of Life Sciences, East China Normal University, Shanghai 200241, China; SymRay Biopharma Inc., Shanghai 200241, China; Acrobiosystems Inc., Beijing 100176, China

**Keywords:** SARS-CoV-2, RBD mutations, ACE2, Neutralizing antibody, IgG

## Abstract

The emergence of coronavirus disease 2019 (COVID-19) pandemic led to an urgent need to develop therapeutic interventions. Among them, neutralizing antibodies play crucial roles for preventing viral infections and contribute to resolution of infection. Here, we describe the generation of antibody libraries from 17 different COVID-19 recovered patients and screening of neutralizing antibodies to SARS-CoV-2. After 3 rounds of panning, 456 positive phage clones were obtained with high affinity to RBD (receptor binding domain). Then the positive clones were sequenced and reconstituted into whole human IgG for epitope binning assays. After that, all 19 IgG were classified into 6 different epitope groups or Bins. Although all these antibodies were shown to have ability to bind RBD, the antibodies in Bin2 have more superiority to inhibit the interaction between spike protein and angiotensin converting enzyme 2 receptor (ACE2). Most importantly, the antibodies from Bin2 can also strongly bind with mutant RBDs (W463R, R408I, N354D, V367F and N354D/D364Y) derived from SARS-CoV-2 strain with increased infectivity, suggesting the great potential of these antibodies in preventing infection of SARS-CoV-2 and its mutations. Furthermore, these neutralizing antibodies strongly restrict the binding of RBD to hACE2 overexpressed 293T cells. Consistently, these antibodies effectively neutralized pseudovirus entry into hACE2 overexpressed 293T cells. In Vero-E6 cells, these antibodies can even block the entry of live SARS-CoV-2 into cells at only 12.5 nM. These results suggest that these neutralizing human antibodies from the patient-derived antibody libraries have the potential to become therapeutic agents against SARS-CoV-2 and its mutants in this global pandemic.

## Introduction

A newly emerged pathogen spreading worldwide named severe acute respiratory syndrome coronavirus 2 (SARS-CoV-2) is now causing a pandemic and is responsible for more than 5,300,000 cases and 340,000 death. This global pandemic threatens public health greatly than ever before. SARS-CoV-2 belongs to *Sarbecovirus* subgenus and shares substantial genetic and functional similarity with other pathogenic human betacoronaviruses, including Severe Acute Respiratory Syndrome Coronavirus (SARS-CoV) and Middle East Respiratory Syndrome Coronavirus (MERS-CoV)^1, 2^. Whereas, the unique pathogenesis and rapid international transmission of SARS-CoV-2 make it become the most infectious and destructive coronavirus than SARS-CoV and MERS-CoV ^3–5^. So, the prophylactic vaccines or therapeutic drugs that are effective and specific to SARS-CoV-2 and its mutants are urgent need to stop the pandemic.

Like other coronaviruses, SARS-CoV-2 utilizes its envelope spike (S) glycoprotein for interaction with a cellular receptor for entry into the target cell. Angiotensin converting enzyme 2 (ACE2) has been identified as the main receptor for viral entry through triggering a cascade of cell membrane fusion events^6^. The spike glycoprotein is composed of two functional subunits responsible for binding to the host cell receptor (S1 subunit) and fusion of the viral and cellular membranes (S2 subunit). And the host range and cellular tropism of SARS-CoV-2 are determinated by the receptor-binding domain (RBD) within the S1 subunit^7^. Although the entry of SARS-CoV-2 into susceptible cells is a complex process and need to be further explored, the receptor-binding and proteolytic processing of the S protein are regarded as the key events in promoting virus-cell fusion. Whereas, more and more RBD mutants have been found under high positive selection pressure during the spread. The fact that these mutations increased the SARS-CoV-2 binding affinity to its host receptor ACE2 reveals a higher risk of more severe infections during a sustained pandemic of COVID-19^8^. Thus, common neutralization of interaction between ACE2 and RBD with different mutations could serve as promising candidates for prophylactic and therapeutic interventions against SARS-CoV-2 infection^9^.

During the last epidemic, SARS patients benefits a lot from convalescent serum collected from recovered subjects^10^. Meanwhile, the blood from COVID-19 patients who have recently become virus-free displayed serum neutralizing activities in a pseudotype entry assay^11^. However, the source of convalescent serum and risk of blood transfusion limited the clinical application. And the heterotransplantation of animal serum may cause anaphylactic reactions such as human anti-mouse antibodies (HAMA) response^12^. Thus, it would be ideal to develop human antibodies against SARS-CoV-2 for prophylaxis or treatment^13^. Different approaches can be developed to get full human antibodies. Among them, phage display library facilitates the identification and development of specific affinity antibodies in a rapid and cost-effective manner. Several full human antibodies against the S1 domain have been generated and all of these antibodies can neutralize the virus *in vitro* and *in vivo*^14^. In order to develop potent neutralizing human antibodies against SARS-CoV-2, single chain antibody fragment (scFv)-phage libraries were constructed by the B cells from 18 different COVID-19 recovered patients. Then several neutralizing human antibodies were identified and characterized with the ability to bind with different RBD and its mutants (R408I, W463R, N354D, V367F and N354D/D364Y) and block the interaction with hACE2. Finally, these antibodies dramatically inhibited SARS-CoV-2 RBD mediated entry into cells using both pseudovirus and live SARS-CoV-2 virus, indicating that patient-derived phage display antibody libraries could be an attractive source of neutralizing antibodies to SARS-CoV-2 and these antibodies are specific prophylactic and therapeutic agents against ongoing SARS-CoV-2 pandemic.

## Results

### Generation of Patient-derived antibody library

As the source of therapeutic antibodies to COVID-19, the anti-SARS-CoV-2 B cells were enriched obvioulsy in COVID-19 convalescent patients. Thus, the PBMC from 18 different COVID-19 recovered patients were isolated to generate phage displayed scFv libraries for panning the neutralizing antibodies (Figure 1A). Prior to construct the phage display library, we examined the binding ability of plasma from 18 recovered COVID-19 patients to SARS-CoV-2 RBD. We found that most of these convalescent patients were able to produce high titer of SARS-CoV-2 RBD-specific antibodies and only three patients mounted relatively lower anti-RBD IgG responses when compared with health donor by enzyme-linked immunosorbent assay (ELISA) (Fig. 1B). Then, the phage display libraries from 17 patients were constructed respectively and quality assessment of each library was undertaken by RT-PCR clone counting and phage display ELISA. The correlation between the size of an antibody library and the likelihood of selecting for the desired antibodies is somewhat intuitive. As shown in Figure 1C, the capacity of primary library from each patient is between 1.01 ×10^7^ to 1.43×10^8^ which is consistent with other immunized scFv library^15^. And the total capacity of whole library is more than 10^9^. Ideally, the size of a library should be equal to its effective size, meaning that all the 10^9^ antibody variants comprising the library are displayed as functional scFv molecules on the phage surface. Whereas, the quality of gene amplification by RT-PCR affects the effective size of the libraries. So, we appraised the positive rate and scFv frequency of each library by PCR and phage display ELISA. Sequence analysis revealed that the framework region (FR) and complementarity determining region (CDR) of selected clones showed the greatest difference in amino acid sequences and the mean positive rate is 95.83% and 88.38% (Figure 1D&E). Taken together, our data revealed that the scFv library from COVID-19 recovered patients were constructed successfully.

**Figure 1.**
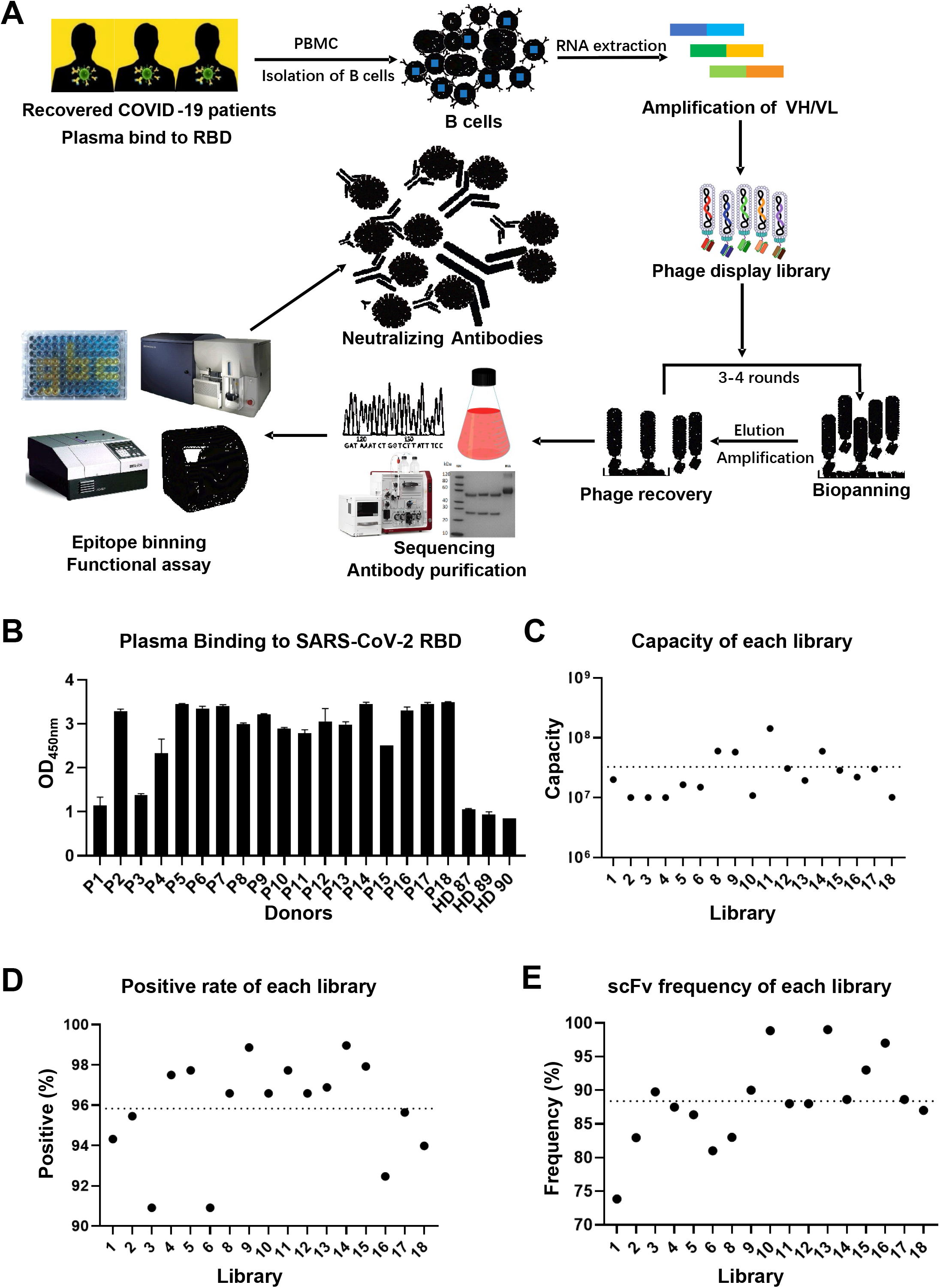
Construction of Patient-derived scFv library. (A) The flow chart of construction and panning of patient-derived scFv libraries. (B) The binding affinity of plasma from each patient to RBD by ELISA. (P1-18, Patients with COVID-19 recovered. HD87-90, healthy donors). (C) The capacity of each library from 17 different COVID-19 recovered patients (7# failed). (D) The clone quality assessment of each scFv library. (E) The scFv display rate of each library.

### Panning against SARS-CoV-2 RBD

Affinity selection of the patient-derived antibody library was performed using solid-phase-bound RBD. After three rounds of panning, 1728 phage clones were randomly picked out to induce the expression of soluble scFv for binding assay by ELISA. As shown in Figure 2A, there are 456 positive clones were identified to bind with RBD specifically. Then, 19 scFv clones with highest affinity to RBD were reconstituted into whole human IgG for functional assay. In order to characterize and group these IgG by the epitope binding regions generated against RBD, the “epitope binning” assay was performed on Octet systems. As shown in Figure 2B, 19 IgG can be divided into 6 different Bins. Among them, Bin1 and Bin2 are biggest groups with 14 and 16 clones suggested that these two epitopes on RBD are most important for recognizing by humoral immune system. So, engineering antibody that targets a specific functioning epitope on an RBD is more important than finding high-affinity, tight-binding mAb, because affinity maturation is a mature and cost-effective technique.

**Figure 2.**
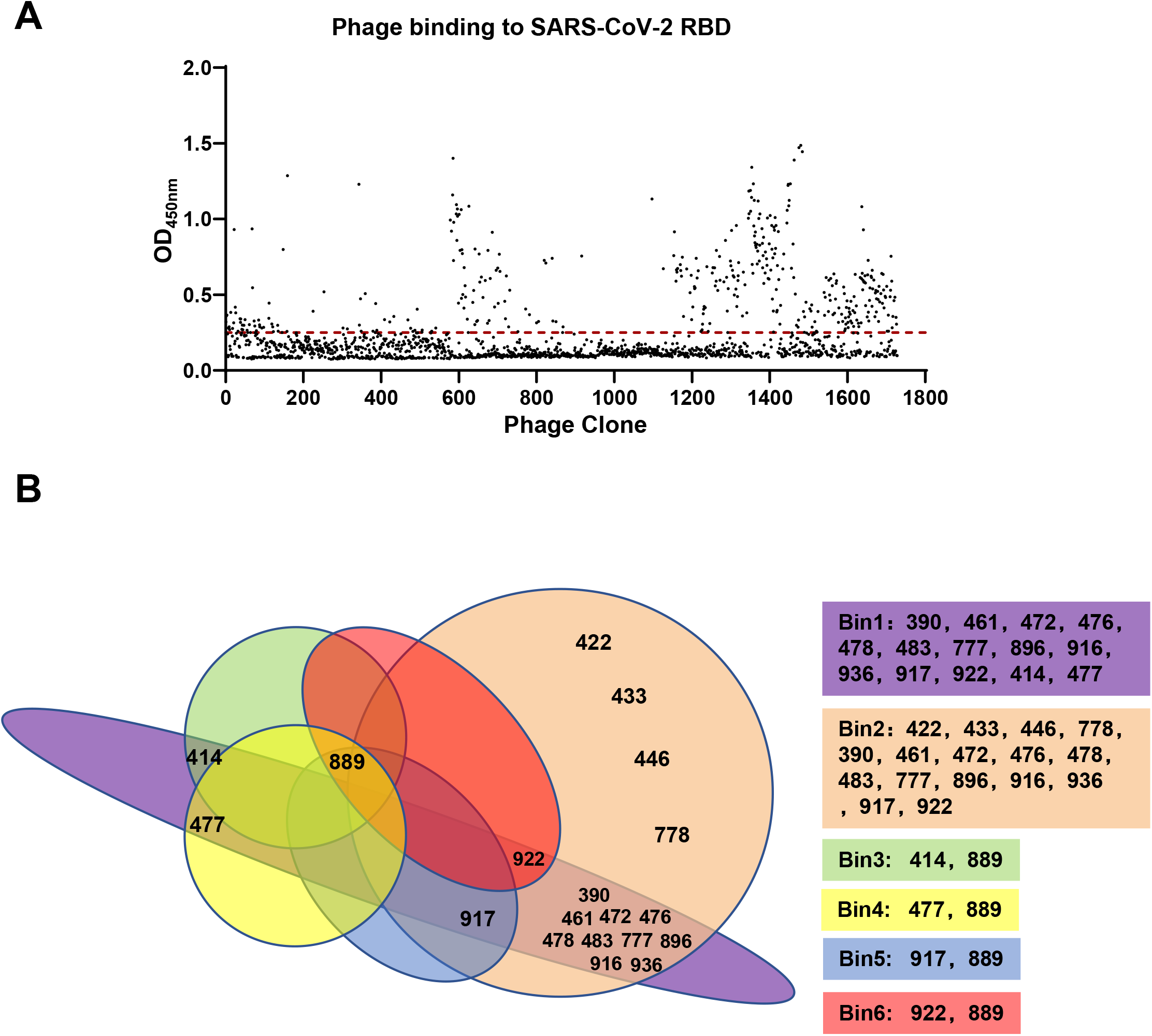
Screening for specific scFv to RBD. (A) The binding affinity of each clone picked after 3 rounds of enrichment. (B) The 19 antibodies were divided into 6 bins by epitope binning assay.

### *In vitro* binding assay to RBD

To further confirm the binding affinity of each IgG in Bin1 and Bin2, the RBD was bound in 96-well plates to do IgG binding assay. Consistent with scFv binding assay (data not shown), the IgG in both Bin1 and Bin2 have similar binding affinity to RBD (Figure 3A&B). As the main step for SARS-CoV-2 entry to the host cells, the interaction between spike and ACE2 is crucial to the viral infection. And the blocking of this interaction by neutralizing antibody have great potential in treating COVID-19. Thus, we sought to investigate whether these IgG can block the binding of RBD to hACE2 overexpressed 293T cells by FACS inhibition assay. As shown in Figure 3C&D, all the IgG in Bin1 and Bin2 have the ability to block the binding of RBD to hACE2 overexpressed 293T cells. Whereas, some IgG in Bin2 group have superior effective blockade of RBD binding to hACE2 cells, suggesting that the epitope of Bin2 is more critical to the infection of SARS-CoV-2 to the host cells. To further confirm this hypothesis, the top 4 IgG in Bin2 were picked out for further study. Although the SARS-CoV-2 is similar to SARS, the amino acid sequence of RBD of SARS-CoV-2 is only about 74% homologous to that of SARS-CoV. So, we used the IgG specific to SARS (CR3022) as a control to do the functional assay. We demonstrated that all the IgG including HTS0422, HTS0433, HTS0446, HTS0483 and CR3022 bind to RBD specifically (Figure 3E). Whereas, only the four IgG (HTS0422, HTS0433, HTS0446 and HTS0483) from Bin2 could inhibit the interaction between RBD and hACE2 (Figure 3F), suggesting that the binding epitope of these IgG is crucial to the SARS-CoV-2 infection.

**Figure 3.**
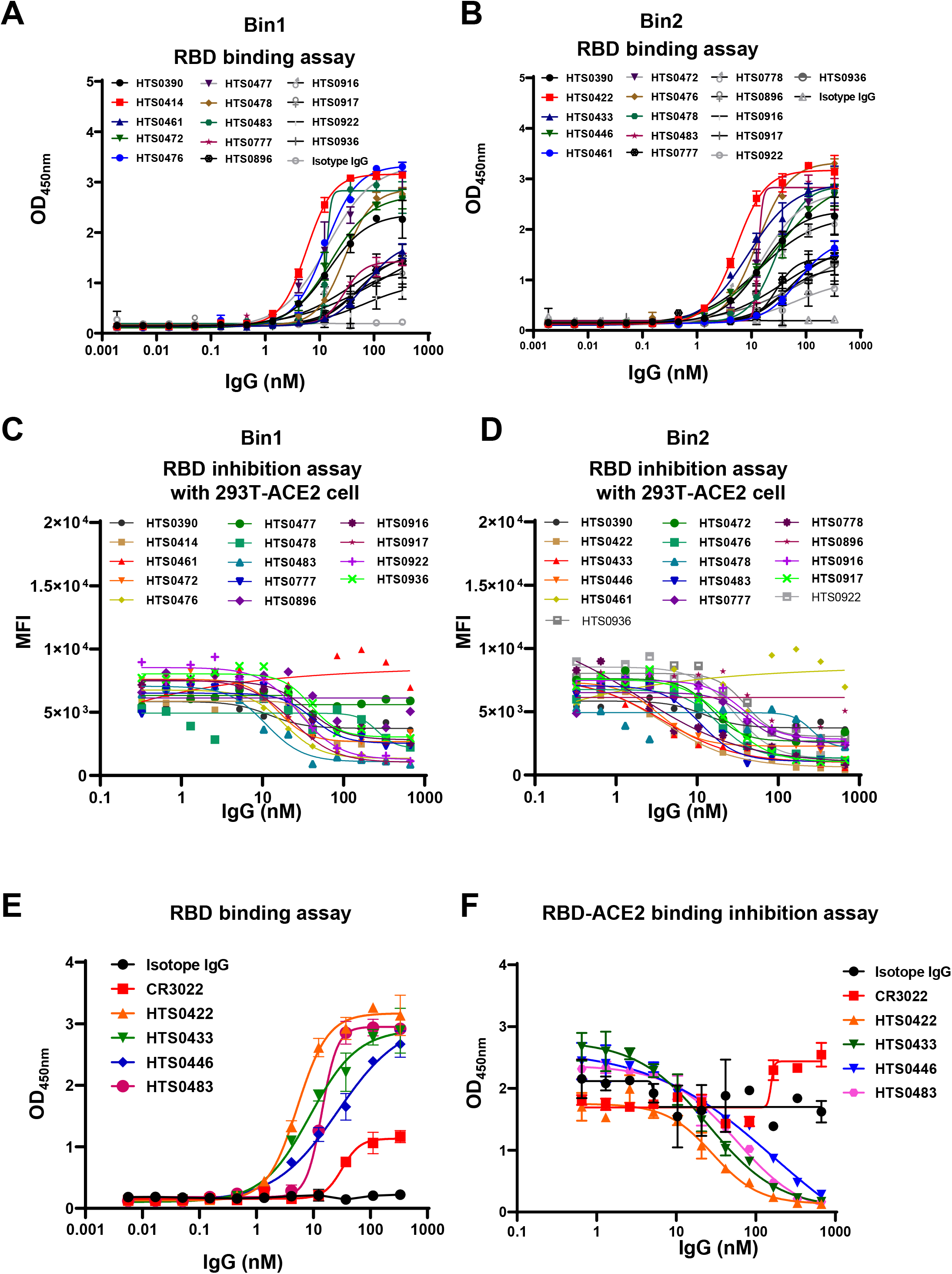
Binding and blocking assays for different scFv. (A, B) Binding of antibodies from Bin1 and Bin2 to RBD protein of SARS-COV-2 is detected by ELISA. (C, D) Blocking the binding of RBD to ACE2 by antibodies from two groups are detected by FACS. Abs were diluted as indicated in the figure. (E) The binding affinity of 4 positive antibodies from Bin2. (F) The blocking ability of 4 positive antibodies from Bin2.

### Cross-neutralization against SARS-CoV-2 RBD mutations

As an RNA virus, mutations in SARS-CoV-2 is commonly found in different countries. Previous studies demonstrated that the mutations in RBD enhanced the binding affinity to the host cells. Therefore, cross-neutralization against SARS-CoV-2 RBD mutations is essential to develop an effective therapeutic antibody. In this study, five different SARS-CoV-2 RBD mutants (R408I, W463R, N354D, V367F and N354D/D364Y) from different strains found in India, Wuhan, Hong Kong, and Shenzhen, respectively, were obtained from ACRO BIOSYSTEMS. As shown in Figure 4A, HTS0422, HTS0433 and HTS0446 bind with both wildtype and mutant (W463R, N354D, V367F, both N354D and D364Y) RBD significantly. Whereas, only HTS0483 bind with R408I, a mutant strain from India, suggesting that R408 is essential for the binding of HTS0422, HTS0433 and HTS0446, and mutations at R408 abolished the binding of antibodies to the virus RBD. And HTS0483 may have broader cross-neutralization effect against SARS-CoV-2 RBD mutations. Accordingly, only HTS0483 inhibit the binding between ACE2 and all RBD mutants including W436R and R408I. Whereas, the antibodies (HTS0422, HTS0433 and HTS0446) can restrict the interaction between ACE2 and wildtype, N354D, V367F or N354D/D364Y (Figure 4B), implying that the mutations at W436R and R408I in RBD may cause immune ignorance by humoral immune response in convalescent patients.

**Figure 4.**
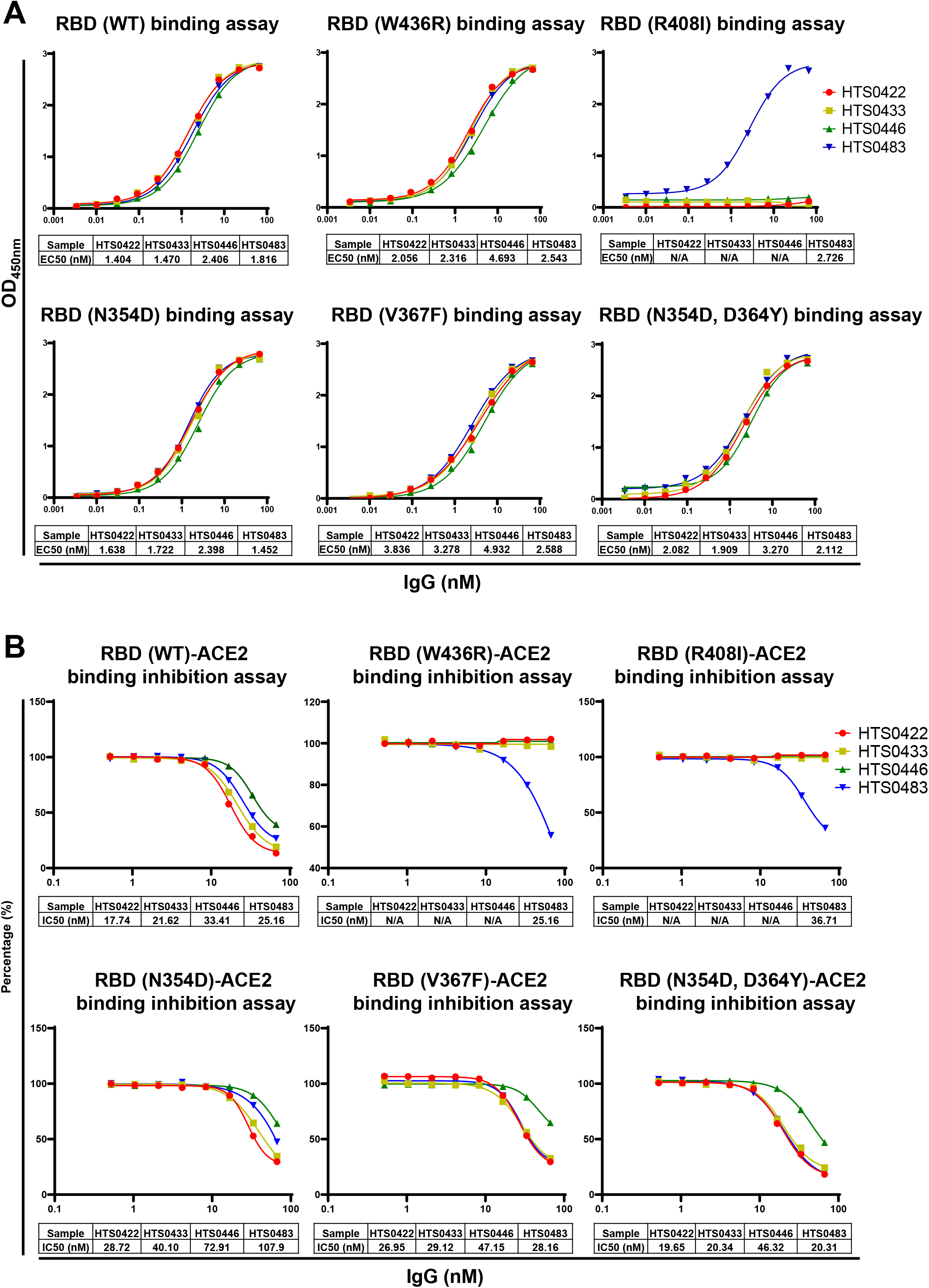
Cross-neutralization against SARS-CoV-2 RBD mutations by different antibodies. (A) The binding affinity of 4 positive antibodies from Bin2 to different SARS-CoV-2 RBD mutations. (B) The blocking ability of 4 positive antibodies from Bin2 to different SARS-CoV-2 RBD mutations.

### Antiviral properties of human antibodies

To explore the potential of these top four IgG as therapeutic drugs, we tested whether these IgG can block the binding of RBD to hACE2 expressed cells by flow cytometry analysis. We found that these top four IgG from Bin2 could restrict the binding of RBD to hACE2-293T cells but not CR3022 which bind to both SARS and SARS-CoV-2 RBD (Figure 5A). Furthermore, we determined the neutralization ability of these four IgG (HTS0422, HTS0433, HTS0446 and HTS0483) using a SARS-CoV-2 S pseudotyped lentiviral particle. Consistent with ELISA and FACS-based blockade result, all these 4 antibodies effectively neutralized pseudovirus entry to host cells ectopically expressing hACE2, with EC50 from 12.80 nM to 16.54nM (Figure 5B&C). Furthermore, SARS-CoV-2 strain hCoV-19/Hangzhou/ZJU-05/2020 was obtained to do the live virus neutralization assay. As shown in Figure 5D, these four IgG from Bin2 inhibit the infection of SARS-CoV-2 significantly at 12.5nM. Finally, authentic infection of Vero-E6 cells with SARS-CoV-2 was neutralized with these IgG antibodies with IC50 from 12.5 nM to 50 nM (Figure 5E), suggesting that these neutralizing antibodies can effectively inhibit infection of the virus to host cells by targeting to the epitope on RBD, and these antibodies are potential therapeutic agents for the treatment of COVID-19.

**Figure 5.**
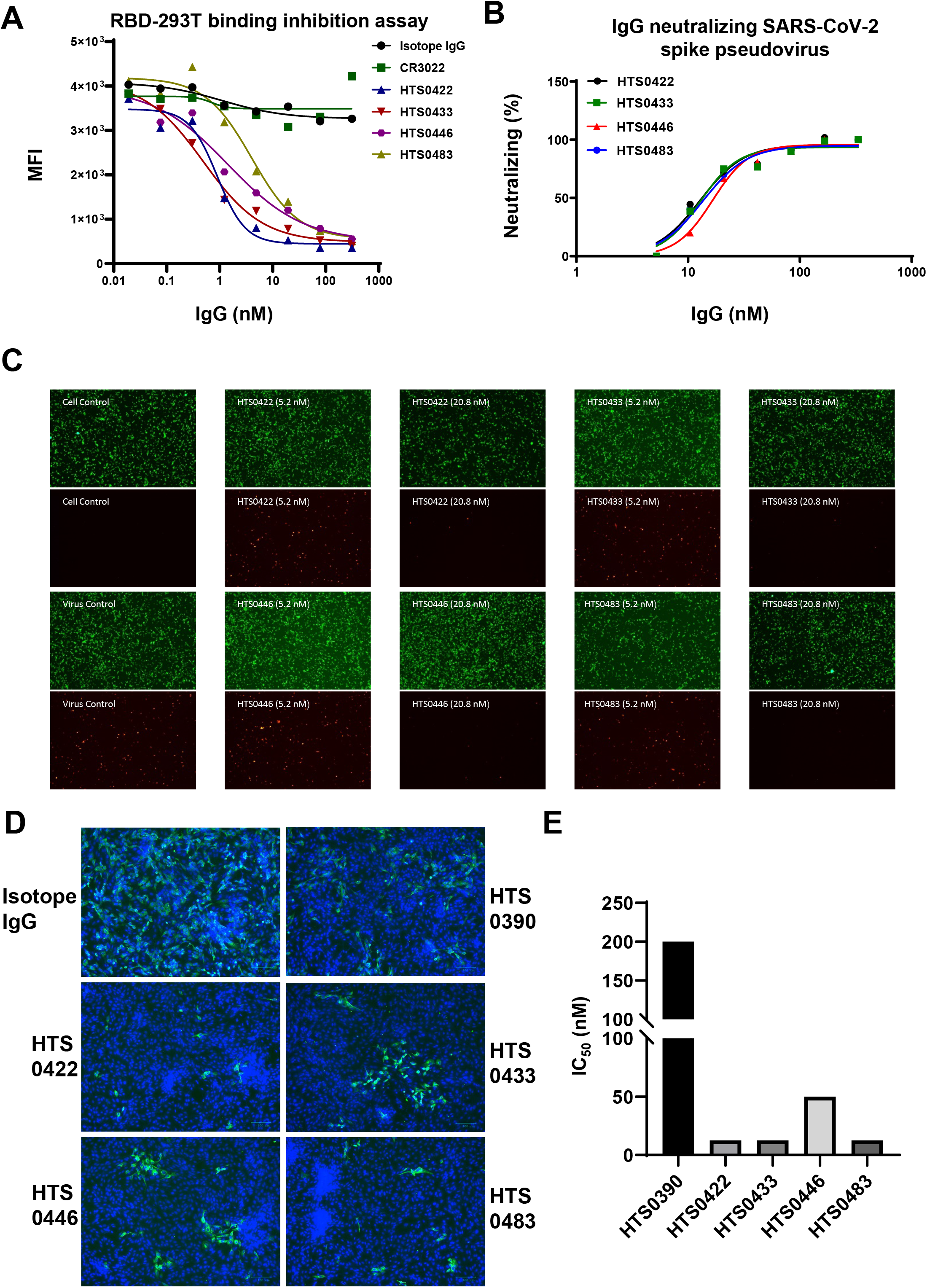
Antiviral properties of human IgG antibodies. (A) The inhibition of 4 positive antibodies from Bin2 to the binding of RBD to hACE2-293T cells. (B) Anti-RBD antibodies (HTS0422, HTS0433, HTS0446, HTS0483) neutralizes viruses pseudotyped with S glycoproteins from the SARS-COV-2 virus. (C) HIVs pseudotyped with the S glycoprotein were incubated with Abs diluted as indicated in the figure for 1 h before infection. Fluorescence intensity in target cells were measured, and the percent neutralization was calculated. Bars indicate SE. (D) The authentic infection of SARS-CoV-2 to Vero-E6 cells was neutralized by Anti-RBD antibodies. (E) The IC_50_ of each antibody from Bin2 was calculated by CPE assay.

## Discussion

Although different approaches have been performed to develop the therapeutic antibodies for infectious diseases, phage display library still persist as the common method for generation of recombinant antibodies against various targets for biomedical applications and research^16^. Till to now, up to 9 fully human therapeutic antibodies discovered via phage display were approved by the Food and Drug Administration (FDA)^17^. In this study, we constructed different phage displayed human scFv libraries from 17 different COVID-19 recovered patients to get the neutralizing antibodies to SARS-CoV-2 infection. Four human antibodies with similar binding epitope were identified to have the potential as a therapeutic antibody for treating COVID-19. Among them, the clone HTS0483 may have broader cross-neutralization against not only wildtype SARS-CoV-2 RBD but also four other mutations.

Similar with other coronaviruses, SARS-CoV-2 use the homotrimeric spike glycoprotein (comprising a S1 subunit and S2 subunit in each spike monomer) on the envelope to bind to their cellular receptors ACE2. Previous study demonstrated that the binding of SARS-CoV and SARS-CoV-2 spike glycoproteins with ACE2 induces the dissociation of the S1 with ACE2, prompting the S2 to transit from a metastable pre-fusion to a more-stable post-fusion state that is essential for membrane fusion^7, 18, 19^. Furthermore, *in vitro* measurements indicate that the RBD is a key functional component within the S1 subunit and is responsible for binding of SARS-CoV-2 by ACE2^20^. Although computer modelling of the interaction between the SARS-CoV-2 RBD and ACE2 has identified some residues that are potentially involved in the interaction, the actual residues involved in the interaction remained unclear. Thus, we cloned the receptor binding domain in S1 subunit of SARS-CoV-2 spike glycoproteins as a target for panning neutralizing antibodies from patient-derived antibody libraries. Although more than 400 positive clones were identified to bind with RBD by ELISA assay, only part of them could block the interaction between RBD and ACE2 significantly, suggesting that not all epitopes in the RBD involved in binding with ACE2. So, we sorted panel of selected antibodies according to epitope specificities by epitope binning assay. The more antibodies that are analyzed for cross-blocking in a pairwise and combinatorial manner against their specific antigen, the higher the probability of discriminating their epitopes. There are 6 different bins were identified, but most antibodies were divided into Bin1 and Bin2. Then we measured the binding affinity and blocking ability of each antibody in Bin1 and Bin2. We demonstrated that the antibodies in Bin1 and Bin2 have similar binding affinity to RBD. However, the neutralizing or blocking ability of antibodies in Bin2 is much better than in Bin1, suggesting that the epitope of Bin2 is essential for ACE2 binding. Therefore, if we can make some point mutation in RBD to explore the binding motif of antibodies in Bin2, the actual residues that mediate the interaction between RBD and ACE2 will be discovered.

Till to now, 32 RBD mutant strains were identified in the strains all around the world and some of them have been proven to increase the infectivity. In addition, the S protein is also important for antigen cognition. Thus, the variation of S protein may change the antigen of virus and influence the vaccine immune efficiency. Therefore, the neutralization antibodies against more SARS-CoV-2 RBD mutations will have better potential in clinical application. In this study, although 4 antibodies are all from Bin2, the binding motif is identical to each other. For example, only HTS0483 binds with R408I mutant in RBD, suggesting that the binding of HTS0483 to RBD is independent of R481. Furthermore, only HTS0483 can block the binding of both W436R and R408I mutants to ACE2 implied that the mutation in RBD may cause the escape of SARS-COV-2 from humoral immune responses. Interestingly, the mutation W436R and R408I are isolated from Wuhan and India respectively. Whereas, the recovered patients are all from Zhejiang province without the mutation in RBD. Therefore, the convalescent patients or vaccine immunized people without the antibody like HTS0483 may still have chance to be infected by the SARS-COV-2 with W436R and R408I mutation. This could be the big challenge to develop effective vaccine against to COVID-19.

Some evidences have shown that the convalescent serum from a patient with a SARS-CoV infection and horse anti-SARS-CoV serum could cross-neutralize the infection of SARS-CoV-2, implied that the structure of RBDs in SARS-CoV-2 and SARS-CoV are similar^2, 6^. Whereas, most antibodies targeted to SARS-CoV have little cross-binding and neutralization activity against spike protein or RBD of SARS-CoV-2, except SARS-CoV antibody CR3022 which binds to the SARS-CoV-2 RBD with a KD of 6.2 nM^21, 22^. In this study, we demonstrated that the binding affinity of CR3022 is lower than 4 antibodies from Bin2. Furthermore, the neutralization activity of CR3022 against RBD of SARS-CoV-2 to ACE2 is vanished when compared with our antibodies, implying that the binding motif of SARS-CoV maybe different from SARS-CoV-2. In addition, the pseudovirus entry and authentic virus infection of Vero-E6 are both inhibited significantly by these antibodies.

In summary, the patient-derived antibody libraries provide another approach for developing therapeutic antibodies against COVID-19 pandemic and the neutralizing monoclonal antibodies will not only have the potential in clinical application but also benefit the mechanism in exploring of SARS-CoV-2 entry to host cells.

## Materials and Methods

### Anti-RBD antibody measurement

18 plasma samples from COVID-19 patients and 3 plasma samples from healthy donors were enrolled in this study. The SARS-CoV-2 antibody ELISA was performed according to the standard ELISA method. Briefly, 30 μl plasma were added to wells coated with recombinant SARS-CoV-2 RBD and incubated for 1 hour at 4°C. Wells were washed three times with PBST (0.1% Tween 20 in PBS) followed by the addition of HRP-conjugated antibody against human IgG and subsequent incubation for 1 hour at room temperature. Wells were washed three times and then TMB was added. Following 20 min of incubation at room temperature, the reaction was stopped and the color reaction was read at 450 nm on an ELISA reader.

### scFv phage display library construction and quality assessment

Total RNA was prepared from the B lymphocytes of convalescent COVID-19 patients, followed by cDNA synthesis. The genes for variable regions of heavy chain and light chain (V_H_ and V_L_) were amplified separately and then cloned into the phage-display vector. The recombinant vector was transformed into Escherichia coli (E. coli) TG1 by electroporation. Cells were plated on bioassay dishes containing 2-YT agar (1.6% tryptone, 1% yeast extract, 0.5% NaCl, 1.5% bacteriological agar) with 100 μg/ml ampicillin and 2% glucose. Following overnight incubation, colonies were scraped off the bioassay dishes and finally stored at −80°C as a primary scFv library. Colony counts of a diluted subpopulation were undertaken to estimate library size. Quality assessment of the library was undertaken by RT-PCR. Amplification of a random selection of 100 clones using the oligonucleotides was undertaken to indicate if clones possessed inserts of the correct size on 1.5% DNA agarose gels. ScFv molecules displayed by rescued phages (M13KO7) were detected via the Myc tag in the scFv-pIII fusion protein using ELISA.

### Selection of phage scFv to SARS-CoV-2

The 96-well plate were coated with RBD in PBS, blocked 2 hours with 2% BSA, and incubated for 3 hours at room temperature with 10^12^ phage scFv in 100 μl PBS containing 2% non-fat milk. After intensive washes with PBST (0.1% Tween 20 in PBS), bound phage antibodies were eluted with 0.1M Glycine-HCl, pH 2.2, and immediately neutralized with 1.0M Tris–Cl, pH 8.8. Eluted phage scFv were subjected to the next round of infection, rescue, and selection. After three rounds of panning, the higher binders to SARS-CoV-2 were selected by ELISA.

### ELISA to identify RBD binders

96-well plates were coated with RBD in PBS, blocked 2 hours with 2% BSA, and incubated with scFv. After three washes with PBST, the bound antibodies were detected by HRP-conjugated anti-Myc antibody followed by incubation with TMD. The color reaction was measured at 450 nm on an ELISA reader.

### Expression and purification of whole IgG antibodies

In order to obtain a large amount antibody, the genes for V_H_ of selected scFv were cloned into pTT5-hIgG1 vector, containing the DNA coding sequence for IgG1-C_H_, and V_L_ cloned into pTT5-hKappa vector separately. The recombinant plasmids were then co-transfected into 293F cells. The antibodies in supernatant were purified by protein G resin.

### Antibody binding ELISA with RBD or RBD mutants

96-well plates were coated with RBD or other RBD mutants in PBS, blocked 2 hours with 2% BSA, and incubated with a serial dilution of purified whole IgG antibodies. After three washes with PBST, the bound antibodies were detected by HRP-conjugated anti-human Fc antibody followed by incubation with TMD and the color reaction was measured.

### Competitive inhibition ELISA assay

RBD or RBD mutants was incubated 1 hour at room temperature with a serial dilution of purified whole IgG antibodies. The above mixture was subjected to 96-well plate coated with ACE2 and incubated for another 1 hour at 4°C. HRP-conjugated anti-mouse Fc was added as secondary antibody. The following procedures were the same as described in ELISA.

### Competitive inhibition FACS assay

293T-ACE2 cells were plated at 1.5×10^5^ cells per well in 96-well plates. The antibodies were diluted in PBS starting at a ratio of 1:4. Serial 4-fold dilutions were mixed with 0.1 μg/ml RBD equally, and incubated at 4°C for 1 hour. The mixture was added to 293T-ACE2 cells, incubated at 4°C for 1 hour, washed 3 times with PBS before APC-conjugated anti-mouse Fc antibody was added. Following 1 hour incubation at 4°C, plates were washed with PBS then developed with FACS.

### BLI assays

Octet systems (Octet RED96e, ForteBio) equipped with amine-reactive, streptavidin, and anti-species sensors were purchased from ForteBio Analytics (Shanghai) Co., Ltd. Epitope binning experiments of anti-SARS-CoV-2 Antibodies were performed in 96-channel mode with in tandem format. HIS1K Biosensors were loaded SARS-CoV-2 RBD protein, His Tag (MALS verified) (Cat. No. S1N-C52H4, ACROBiosystems). Then interact with the first antibody and the second antibody in sequence, and detect the binding signal of the second antibody to determine whether the two antibodies recognize the same epitope. The in tandem style assay comprised a five-step binding cycle; 1) a buffer baseline was established for 1 min, 2) 5 μg/ml SARS-CoV-2 RBD protein was captured about 0.4nm, 3) 7.5-15 μg/ml mAb array was loaded to saturate the immobilized antigen for 1 min, 4) 15 μg/ml of the test mAb was bound for 1min, and 5) the capture surfaces were regenerated for 30 sec. HIS1K biosensors were regenerated with 10 mM Glycine-HCl, pH1.5 for 5 sec with 3 times and neutralized for 5 sec with 3 times in neutralization buffer immediately after each regeneration. In tandem assays were conducted depending on the experiment.

### Preparation of RFP/SARS-CoV-2 spike pseudovirus

Production of VSV pseudotyped with SARS-CoV-2 spike was performed using the standard lentivirus production method. Briefly, 293T cells were transfected with psPAX2 and vectors encoding SARS-CoV-2 spike protein as well as a core plasmid expressing RFP. After 48 hours post-infection, RFP/SARS-CoV-2 spike pseudoviruses were packaged and collected from the culture supernatant.

### Neutralization activity of mAbs against pseudovirus

RFP/SARS-CoV-2 spike pseudovirus system was used as an infection model to evaluate the neutralization activity of antibodies. First, the RFP/SARS-CoV-2 spike pseudoviruses were pre-incubated with a serial dilution of purified whole IgG antibodies. Then, 293T-ACE2 cells were infected by the RFP/SARS-CoV-2 spike pseudovirus with or without antibodies. After 2 hours incubation at 37°C, the medium was refreshed with DMEM containing 10% FBS and continuously cultured for another 72 hours. The infected 293T-ACE2 cells were observed under microscope and the neutralizing IC50 was calculated after FACS detection.

### Neutralization activity of mAbs against live SARS-CoV-2

SARS-CoV-2 obtained from a sputum sample was amplified in Vero-E6 to make working stocks of the virus. To analyze the mAb neutralizing activities, two-fold serial dilutions of mAbs were added to the same volume of 100 TCID50 of SARS-CoV-2 and incubated for 1 hour at 37°C. The mixture was added to a monolayer of Vero-E6 cells in a 96-well plate and incubated at 37°C. Cytopathic effects (CPE) were observed and recorded from day 4 to day 6. For immunofluorescence (IF) analysis, Vero-E6 cells were transfected and fixed on day 6 post-transfection in 80% acetone for 10 min at room temperature. Cells were immunolabelled for 1.5 hours at room temperature with the rabbit anti SARS-CoV-2 spike antibody, washed three times with PBS, and followed by the addition of Alexa Fluor 488-conjugated goat anti-rabbit IgG antibody. Wells were washed three times and then DAPI was added. Following 30 minutes of incubation at room temperature, the reaction was reading in fluorescent microscopy.

## Acknowledgements

This work was supported by National Key R&D Program of China [2019YFA0802800, 2018YFA0507001]; Zhejiang University special scientific research fund for COVID-19 prevention and control (2020XGZX086); Zhejiang Provincial Science and technology department key R&D plan emergency project [2020c03123-8]; Innovation Program of Shanghai Municipal Education Commission [2017-01-07-00-05-E00011]; Fundamental Research Funds for the Central Universities

